# The two-step chemosensory system underlying the oligophagy of silkworm larvae

**DOI:** 10.1101/719344

**Authors:** Kana Tsuneto, Haruka Endo, Fumika Shii, Ken Sasaki, Shinji Nagata, Ryoichi Sato

**Affiliations:** Graduate School of Bio-Applications and Systems Engineering, Tokyo University of Agriculture and Technology, Koganei, Tokyo 184-8588, Japan; Department of Integrated Bioscience, Graduate School of Frontier Sciences, The University of Tokyo, Kashiwa, Chiba 277-8562, Japan; Graduate School of Agriculture, Tamagawa University, Machida, Tokyo 194-8610, Japan

**Author notes:** Correspondence (H.E.) or (R.S.).

## Abstract

Oligophagous insect herbivores specifically identify host-plant leaves using their keen sense of taste^1^. Plant secondary metabolites and sugars are key chemical cues for insects to identify host plants and evaluate their nutritional value, respectively^2^. However, it is poorly understood how the insect chemosensory system integrates the information from various gustatory inputs. Here we report that a two-step chemosensory system is responsible for host acceptance by larvae of the silkworm *Bombyx mori*, a specialist for several mulberry species. The first step controlled by a gustatory organ, the maxillary palp (MP), is host-plant recognition during palpation at the leaf edge. Surprisingly, MP detects chlorogenic acid, quercetin glycosides, and β-sitosterol, which stimulate feeding by the silkworm^3–6^, with ultra-sensitivity (thresholds of aM to fM). Detecting a mixture of these compounds triggers test biting. The second step is evaluation of the sugar content in the leaf sap exuded by test biting. Low-sensitivity chemosensory neurons in another gustatory organ, the maxillary galea (MG), mainly detect sucrose in the leaf sap exuded by test biting, allowing larvae to accept the leaf and proceed to persistent biting. Our present work shows the behavioral and neuronal basis of host acceptance in the silkworm, mainly driven by six phytochemicals. It also reveals that the ultra-sensitive gustation via MP strictly limits initiation of feeding in the silkworm unless it detects a certain combination of host compounds, suggesting an essential role of MP in host-plant selection. The two-step chemosensory system reported herein may commonly underlie stereotyped feeding behavior in phytophagous insects and determine their feeding habits.

## Main

Host-plant selection by phytophagous insects is dependent on their acceptance or rejection of plants. To clarify the mechanism of host-plant acceptance by silkworm larvae, we observed larval feeding behavior towards a host leaf from white mulberry *Morus alba*. When a silkworm encounters a leaf, it first palpates the leaf edge using a peripheral gustatory organ known as the maxilla, intermittently bites the edge several times, and finally engages in continuous biting (2–3 times per second) with its head shaking in the dorso-ventral direction along the leaf edge (Fig. 1a, Supplementary Video 1). The intermittent biting with palpation and the continuous biting with head-moving are termed test biting and persistent biting, respectively^7,8^. We hypothesized that sensing of chemical cues from a *M*. *alba* leaf via the maxilla induces test biting because test biting always occurs after palpation with the maxilla. The maxilla consists of the maxillary palp (MP) and maxillary galea (MG) (Fig. 1b). To assess the roles of MP and MG in induction of test biting, we used MP- or MG-ablated larvae. MP-ablated larvae showed palpation, but no test or persistent biting (Fig. 1c, d, Supplementary Video 2). MG-ablated larvae showed palpation and test biting, and stopped biting within 1 minute, and did not progress to persistent biting (Fig. 1c, d, Supplementary Videos 3). When we ablated an olfactory organ antenna (AN), the larvae showed palpation, test biting, and persistent biting similar to intact larvae (Fig. 1c, d, Supplementary Video 4). These results indicate that MP and MG are essential for induction of test and persistent biting, respectively.

**Figure 1.**
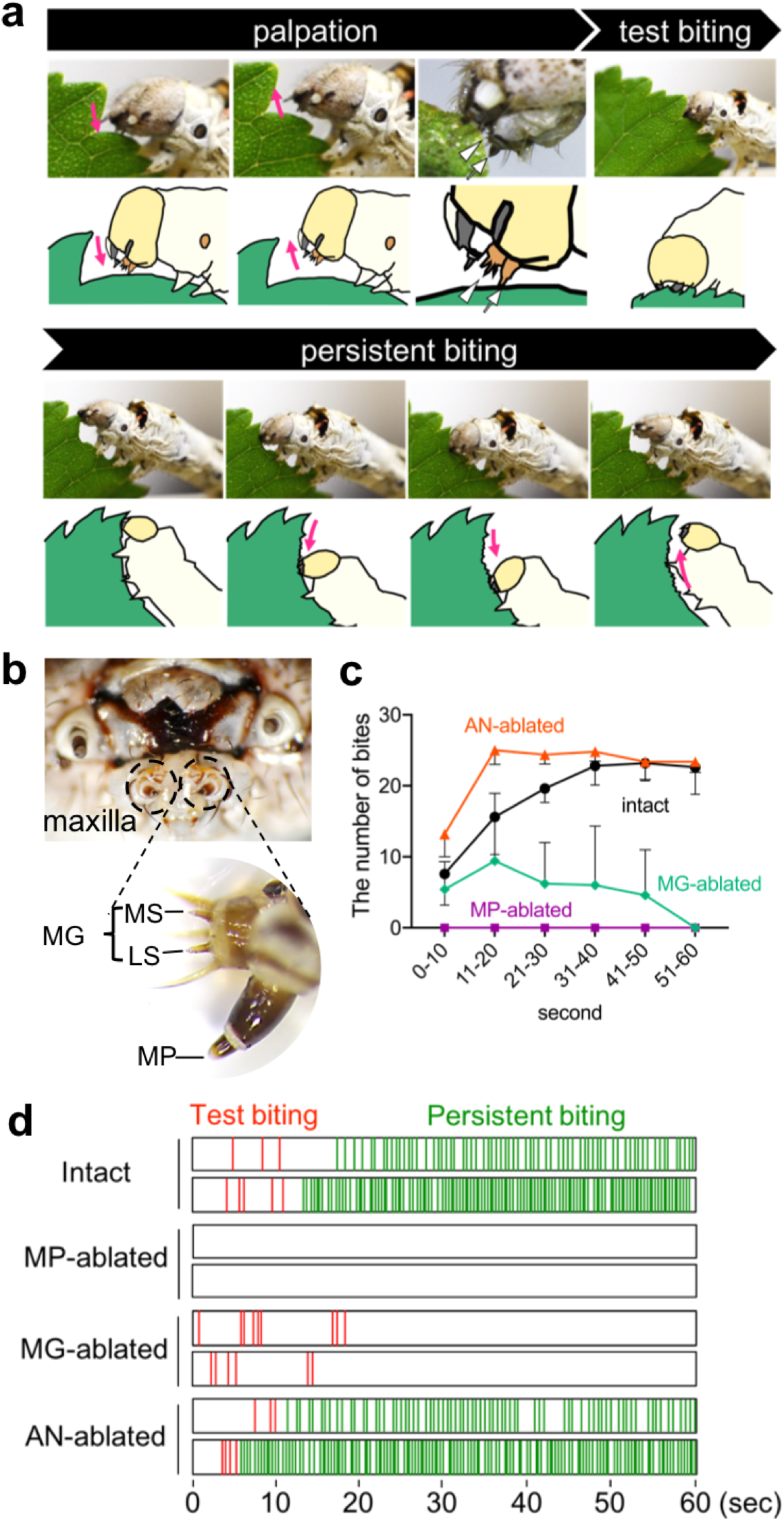
Peripheral gustatory organs, maxillary palp (MP), and maxillary galea (MG) are responsible for test and persistent biting, respectively, of a mulberry leaf. **a,** Feeding on mulberry leaves by silkworm larvae: (1) Palpation; a silkworm larva first palpates the leaf-surface with its maxilla (MP and MG) for 5-30 seconds. The white arrow and arrowhead indicate MP and MG, respectively. (2) Test biting; the larva bites the leaf edge several times intermittently during palpation. (3) Persistent biting; the larva nibbles the leaf edge repeatedly (2-3 times per second) with its head moving in the dorso-ventral direction along the leaf edge. magenta arrows indicate the direction of head movement. **b,** Mouthparts of a silkworm larva. Upper, larval mouthparts including one pair of maxilla. Lower, higher-magnification view of the maxilla. The maxilla consists of the MG and MP. The MG has two gustatory sensilla, the lateral and medial styloconic sensilla (LS and MS). **c,** Frequency of biting by intact, MP-ablated, MG-ablated, and antenna (AN)-ablated larvae of mulberry leaves over a 10-second period. Data are means ± SD (n = 5). **d,** Representative raster plots of the timing and duration of the biting behavior of larvae in **c.** Red and green lines indicate test and persistent bites, respectively.

To assess whether such MP- and MG-controlled biting contributes to the oligophagy of the silkworm, we observed feeding behavior towards leaves of various plant species. The silkworm is a specialist for some *Morus* species, including *M. alba*. In addition, the leaves of Cichorioideae plants of the Asteraceae (*e.g*., dandelion [*Taraxacum officinale*] and Indian lettuce [*Lactuca indica*]) are consumed a relatively small amount by silkworm larvae^9, 10^. Of the larvae, 88.9 ± 5.3% showed test biting of *M. alba* within 1 minute after reaching the leaf edge, compared to 70.0 ± 10.0 and 63.3 ± 13.3% for *Sonchus oleraceus* and *T*. *officinale*, respectively (Fig. 2a). Of the larvae, 80 ± 11.5, 53.3 ± 14.5, and 30 ± 10% proceeded to persistent biting of *M*. *alba, S. oleraceus,* and *T*. *officinale*, respectively. In contrast, the larvae had a lower probability of test biting (3–33%) of 12 inedible leaves, which were finally rejected by most of them (Fig. 2a, b). Thus, the probability of test biting was higher for edible than for inedible leaves and persistent biting was induced only by edible leaves. These results suggest that the host plant is recognized prior to biting.

**Figure 2.**
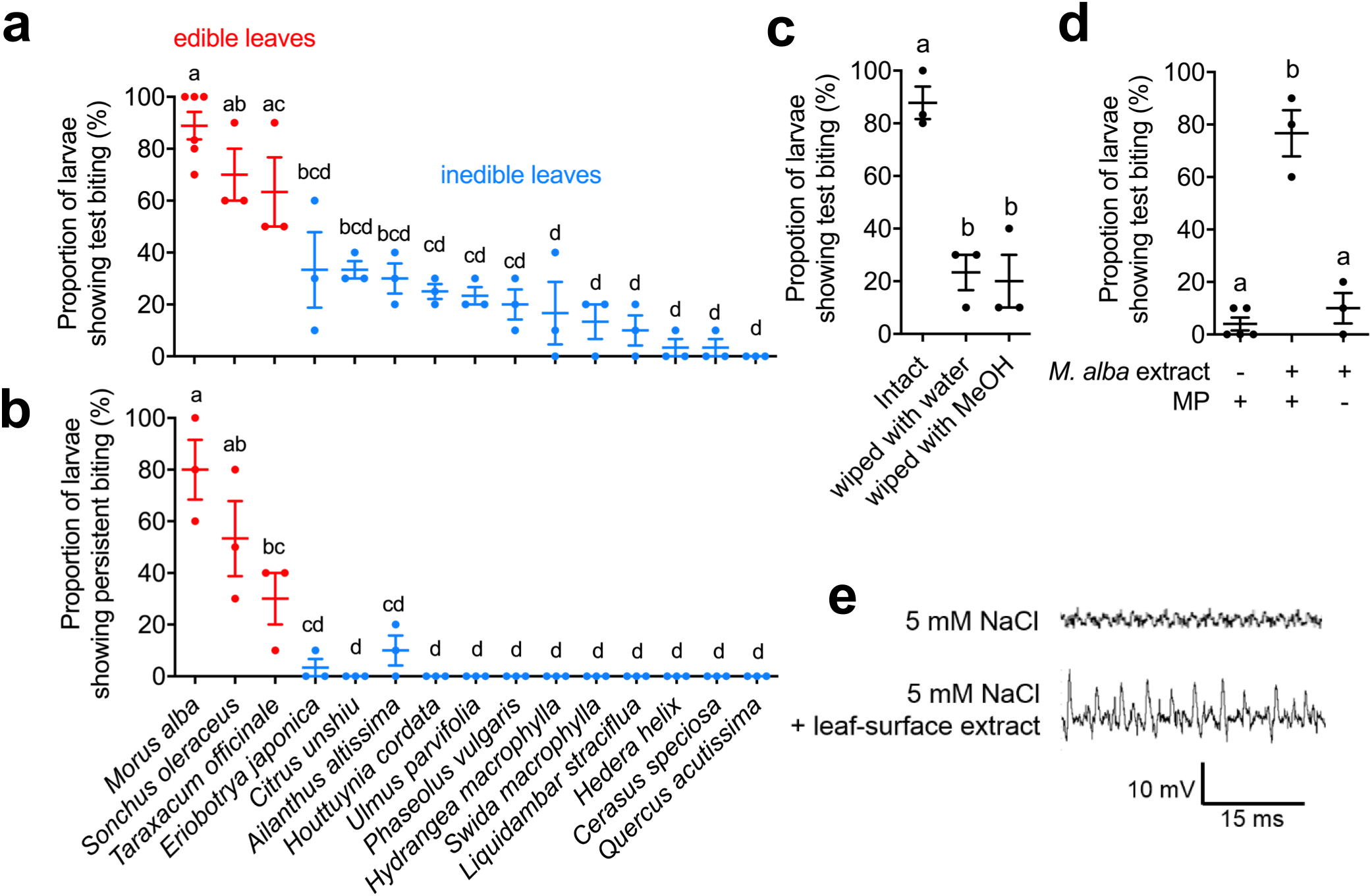
The MP detects leaf-surface compounds and triggers test biting of edible leaves. **a, b,** Proportion of larvae showing test biting **(a)** and persistent biting **(b)** of edible and inedible leaves. **c, d,** Proportion of larvae showing test biting of intact mulberry leaves or mulberry leaves wiped with sheets of Kimwipe® paper moistened with water or methanol **(c)** or filter paper treated with methanol or mulberry leaf-surface extract **(d)**. MP + and MP − indicate intact and MP-ablated larvae, respectively. **a-d,** Biting of 10 larvae was observed for 1 minute. Experiments were repeated as independent biological replicates (n = 3−5). Data are means ± SE. The same letters indicate no significant difference (*P* > 0.05) by one-way analysis of variance (ANOVA) followed by Tukey *post hoc* test. **e,** Typical spikes from chemosensory sensilla of the MP towards 5 mM NaCl (negative control) and leaf-surface extract (Compounds from a leaf/10 ml).

Insects sense leaf-surface compounds during palpation^1^. To elucidate whether leaf-surface compounds induce test biting, we wiped *M. alba* leaves with water or methanol, which markedly decreased the proportion of larvae showing test biting (Fig. 2c). Conversely, 76.7 ± 8.8%, 37.5 ± 10.3%, and 36.0 ± 10.8% of the larvae showed test biting of filter paper treated with methanol and leaf-surface extracts of *M*. *alba*, *S*. *oleraceus*, and *T. officinale*, respectively (Fig. 2d, Extended Data Fig. 1a-c). MP ablation diminished test biting towards filter paper treated with an extract of *M*. *alba* leaf-surface (Fig. 2d). Furthermore, tip recording of the sensilla in the MP^12^ revealed that MP neurons responded to the leaf-surface extract (Fig. 2e). These findings indicate that compounds in edible leaf-surface extracts stimulate the MP and trigger test biting.

To identify inducers of test biting, we searched for secondary metabolites in edible leaves of *M*. *alba*, *Lactuca indica*, and *T*. *officinale* in the plant-metabolite database KNApSAcK^11^ because secondary metabolites are thought to be key chemical cues for host-plant recognition^2^. The search yielded chlorogenic acid (CGA) and quercetin-3-*O*-rhamnoside (Q3R). CGA reportedly increases the amount of food intake by the silkworm^6^: Q3R is an analog of isoquercitrin (ISQ), which reportedly induces biting by the silkworm^5^. In addition, we focused on β-sitosterol (βsito) because it also reportedly induces biting by the silkworm^4^. We first recorded the responses of MP and MG towards these compounds. Surprisingly, MP responded to the four compounds at the attomolar and femtomolar levels (Fig. 3a). Based on the shape, amplitude, and frequency of the spikes, CGA, ISQ, Q3R, and βsito were estimated to stimulate at least one, three, two, and one neuron(s), respectively. It is plausible that this ultra-sensitivity of the MP enables detection of trace amounts of CGA, Q3R, ISQ, and βsito at the leaf-surface. In contrast, the MG did not respond to these compounds (Extended Data Fig. 2b, 2c). Next, we assessed whether the four compounds induce test biting. Filter paper, which were treated with each single compound, mixtures of two compounds, and a mixture of ISQ, Q3R, and βsito induced test biting by 20–40% of larvae. In contrast, mixtures of three compounds (CGA+ISQ+βsito, and CGA+Q3R+βsito) and the mixture of all four compounds resulted in a high probability of test biting comparable to the *M*. *alba* leaf-surface extract (Fig. 2d and 3b), but did not induce persistent biting (Extended Data Fig. 1d). Filter papers treated with extremely dilute mixtures of CGA, ISQ, and βsito still induced test biting to some extent (Fig. 3c). Meanwhile, a mixture of fructose, sucrose, glucose, and *myo*-inositol did not induce biting (Fig. 3b). These results suggest that a trace amount of set of CGA+ISQ/Q3R+βsito contribute to host recognition and induction of test biting.

**Figure 3.**
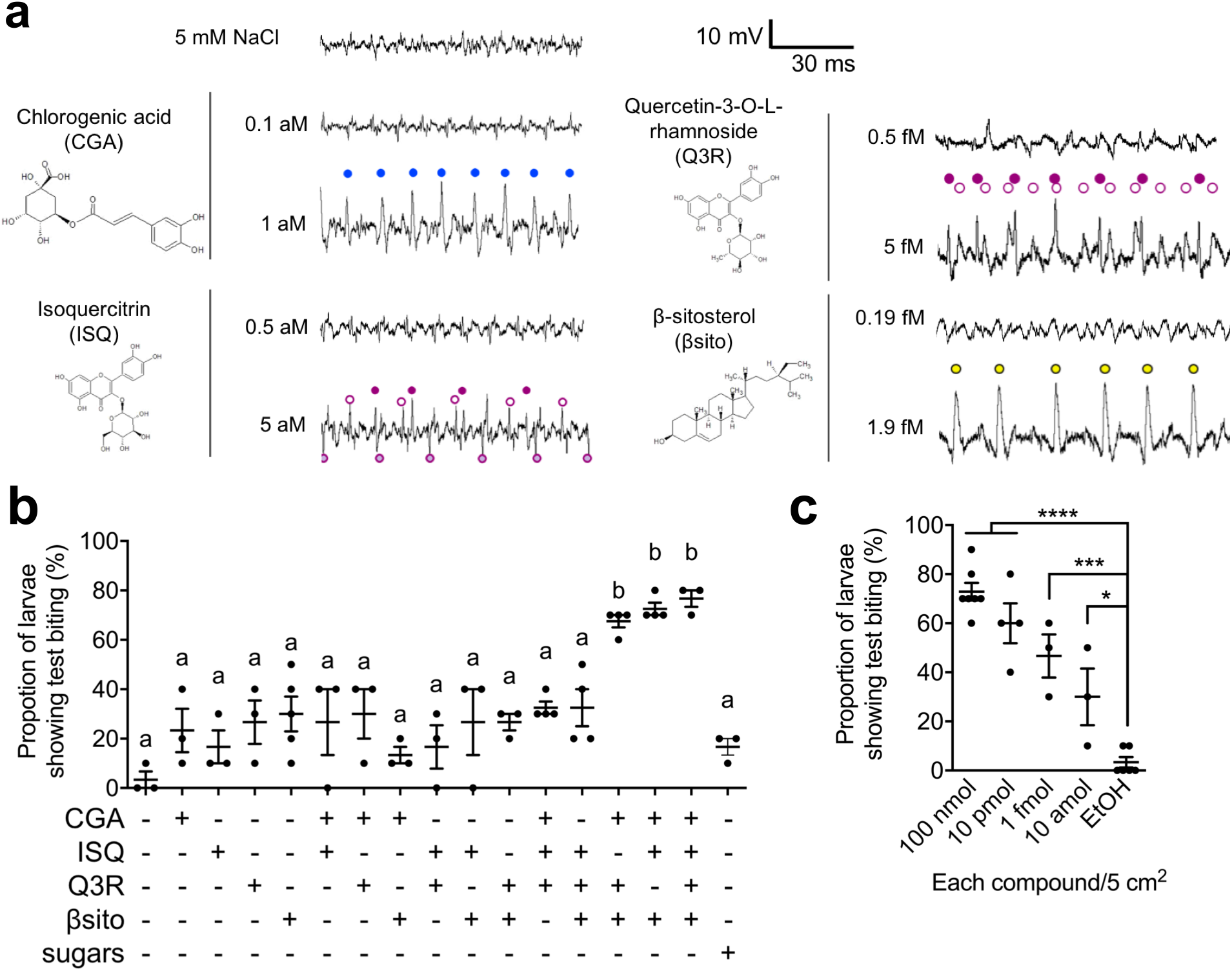
A mixture of chlorogenic acid, isoquercitrin/quercetin-3-*O*-rhamnoside, and f3-sitosterol induces test biting by stimulating ultra-sensitive MP sensory neurons. **a,** Typical electrophysiological recordings from the MP in response to chlorogenic acid (CGA), isoquercitrin (ISQ), quercetin-3-*O*-rhamnoside (Q3R), and β-sitosterol. **b, c,** Fraction of larvae showing test biting of filter paper treated with CGA, ISQ, Q3R, βsito, and sugars (sucrose, fructose, myo-inositol, and glucose) **(b),** and with an extremely dilute mixture of CGA, ISQ, and βsito **(c).** Filter paper was treated with 100 μL of 1 mM stimulant solution (100 nmol of each compound/5 cm^2^) **(b)** or a dilution series of the CGA, ISQ, and βsito mixture **(c).** Biting by 10 larvae was observed for 1 minute. Experiments were repeated as independent biological replicates (n = 3−5). Data are means ± SE. Statistical analysis was performed using one-way analysis of variance (ANOVA) followed by Tukey *post hoc* test. The same letters indicate no significant difference (P > 0.05). An asterisk indicates a significant difference (\**P* < 0.05; \*\*\**P* < 0.001; \*\*\*\**P* < 0.0001).

Next, we investigated the role of MG in inducing persistent biting. The lateral sensillum (LS) in the MG is involved in recognition of feeding stimulants. The LS has three neurons specifically tuned to glucose, sucrose, and *myo*-inositol at around the millimolar level^14^. Therefore, we hypothesized that sugars in the leaf sap exuded by test biting stimulate the MG and induce persistent biting. As feeding initiation is strictly regulated by test biting, we conducted an agar-based food intake assay using starved larvae^16^ to simply evaluate persistent biting. In this assay, starved larvae no longer exert strict oligophagy and sometimes randomly bite; this biting substitutes for test biting, and consequently persistent biting occurred in the absence of the inducers of test biting. Alternatively, unlike when feeding on leaves, the MG directly detected high concentrations of compounds at the agar surface during palpation, resulting in induction of persistent biting. Larvae fed agar diet containing sucrose at > 10 mM with showing persistent biting (Fig. 4a). The amount of food intake seemed to correlate with the length of persistent biting. Indeed, a sucrose dose-dependent increase in the body weight was observed in intact and MP-ablated larvae (Extended Data Fig. 3b, c). The magnitude of the sucrose-induced increase in larval weight was significantly smaller in MG-ablated larvae than in intact larvae (Fig. 4a), suggesting an important role for MG in modulating the amount of food intake. Meanwhile, *myo-*inositol and glucose themselves did not induce larval weight increase (Fig. 4b and Extended Data Fig. 3d), whereas *myo*-inositol showed a supplemental effect in the presence of sucrose (Fig. 4b) in consistent with a previous study^16^. These results suggest that sucrose and *myo*-inositol contribute to induction of persistent biting by stimulating MG.

**Figure 4.**
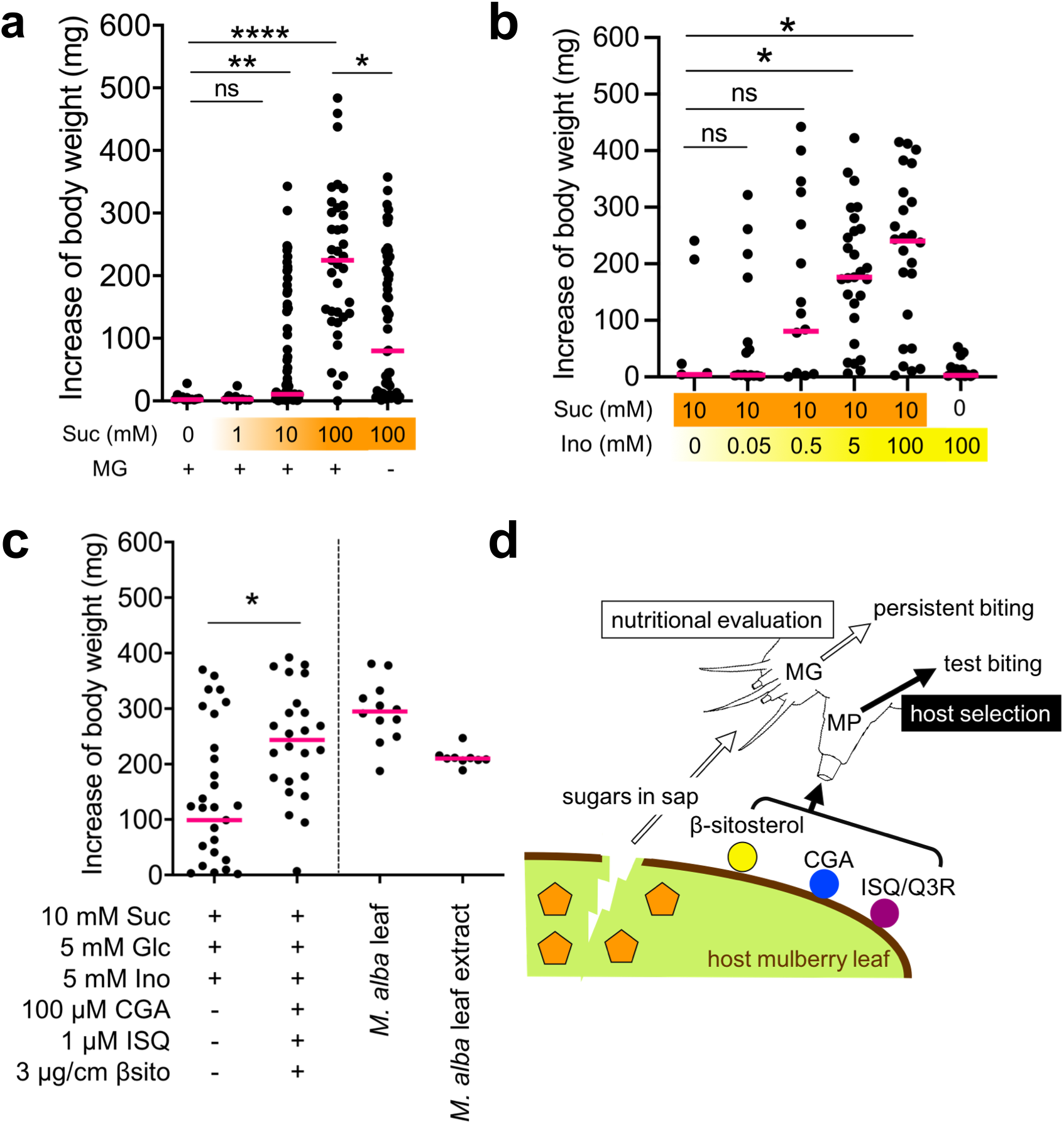
Sucrose induces persistent biting and food intake, which are modulated by *myo*-inositol and inducers of test biting. **a, b,** Effects of sucrose **(a),** *myo*-inositol **(b),** and inducers of test biting **(c)** on the increase in larval body weight by agar-based food intake assay. b, Effect of ablation of the MP and MG on the sucrose-dependent increase in larval weight. The agar-based diet contained 100 mM sucrose. **a-c,** Fifth-instar larvae were starved for 3 days beginning at the end of molting. The body weight of the three larvae increased after feeding of the agar-based diet (9% cellulose and 1% agar) for 3 hours. Magenta bars denote median. Statistical analysis was performed using Kruskal-Wallis test followed by Dunn test (**a, b**) and Mann-Whitney *U*-test (**c**). "ns" indicates no significant difference; an asterisk indicates a significant difference (\**P* < 0.05; \*\**P* < 0.01; \*\*\*\**P* < 0.0001). MG + and MG − indicate intact and MG-ablated larvae, respectively. **d,** Proposed model of host recognition and acceptance by the silkworm.

Finally, we assessed whether that the inducers of test biting (CGA, ISQ, and βsito) and sugars (sucrose, *myo*-inositol, and glucose) resulted in an increase in larval weight similar to that induced by *M. alba* leaf extract and intact leaf. A mixture of 10 mM sucrose, 5 mM *myo*-inositol and 5 mM glucose, similar to the concentrations in *M. alba* leaves^17^, resulted in an increase in larval weight in the presence, but not the absence, of a mixture of sugars, similar to a *M. alba* leaf and a leaf-extract-containing agar-based diet (Fig. 4c). The test biting induced by CGA+ISQ+βsito accelerated the first bite (Extended Data Fig. 4), which might cause persistent biting and consequently increased the total food intake. Therefore, we concluded that these six compounds are major phytochemical drivers of silkworm feeding (Fig. 4d).

Since the 1970s, entomologists have noticed that insects identify their host plants at the leaf-surface, but the mechanism remains unknown. We report here the ultra-sensitive gustation by MP underlies host-plant recognition at leaf-surface in the silkworm. CGA, ISQ/Q3R, and βsito are not specific to edible leaves and other phytochemicals may be also involved in host-plant recognition. Nevertheless, it is plausible that the presence of all the three compounds is a marker of mulberry leaves at least in the ecosystem in which *Bombyx madarina*, an ancestor of the domesticated silkworm, resides. Additionally, rejection of inedible leaves upon detection of feeding deterrents^13,19^ seems to facilitate strict host plant selection. Host-plant recognition by MP may be conserved among phytophagous insects. For example, insect oligophagy and polyphagy are likely consequences of strict and loose restriction, respectively, of feeding initiation by MP. Tuning of neurons in MP for unique combinations of phytochemicals must reflect insect-plant relationships.

## Acknowledgements

We thank T. Shimada and H. Takai (University of Tokyo) for technical assistance in electrophysiological experiments; N. Watanabe (Tokyo University of Agriculture and Technology) for assistance in sampling various plants leaves; Y. Banno (Kyushu University) for providing fresh *M. alba* leaves. This research was financially supported by Japan Society for the Promotion of Science (JSPS) KAKENHI (Grant Number 17K19261 and 18J00733) to R.S. and H.E.

## Author contributions

K.T., H.E. and R.S. conceived and designed the study. K.T. performed most experiments. H.E. contributed to analysis of leaf surface compounds under the supervision by S.N. F.S. performed some behavioral experiments. K.S. supervised electrophysiological experiments. K.T., H.E. and R.S. wrote the manuscript with intellectual inputs from all the authors. H.E. and R.S. supervised the study.

## Materials & Correspondence

Correspondence and requests for materials should be addressed to H.E. or R.S.

## Methods

### Insects

The silkworm (*B. mori* Kinshu × Showa hybrid) eggs were purchased from Ueda Sanshu Ltd (Nagano, Japan). The silkworms were reared on an artificial diet, Silkmate 2M (Nihon-Nosan Co. Ltd., Kanagawa, Japan) with 16L-8D at 25°C. Larvae were provided with fresh diets every day to synchronize growth.

### Biting assay towards leaves and filter paper

Videos of feeding behavior towards leaves and filter paper were taken by the D5100 camera (Nikon, Tokyo, Japan) equipped with a telephoto lens AF-S DX Micro Nikkor 85mm (Nikin, Tokyo, Japan). Larvae were used 24 to 48-h after moulting to the fifth instar. To prevent non-specific biting due to starvation, all larvae fed artificial diets for two minutes before use. For elucidation of the role of chemosensory organs, maxillary palp, maxillary galea or antenna was quickly removed from a larva with fine forceps after the two-minute feeding. Videos were taken for around 30 to 60 seconds after starting palpation towards leaves and filter papers. Fresh host and non-host leaves were collected within 2 weeks before use around the campuses of Tokyo University of Agriculture and Technology (Fuchu and Koganei, Tokyo, Japan) and stored in 4°C. For preparation of wiped leaves and leaf-surface extract, fresh leaves were gently wiped with kimwipes (Nippon Paper Crecia Co., Tokyo, Japan) wet with water or methanol. Kimwipes were dried in a drying machine, wet with water or methanol again and put into a centrifugal filter column (Merck Millipore, Darmstadt, Germany), and centrifuged to collect leaf surface extracts. Filter paper (size: 1 cm long, 5 cm wide) was treated with surface extracts from a leaf. A larva moved freely after being put on a leaf and all larvae started finally palpation toward all kinds of plant leaves. For biting assay using commercial compounds, filter paper was impregnated with 100 μl stimulant solution. Chlorogenic acid (Nacalai tesque, Kyoto, Japan), isoquercitrin (Extrasynthese, Lyon, France), quercetin-3-*O*-rhamnoside (Extrasynthese) and/or β-sitosterol (Abcam, Cambridge, UK) were dissolved in ethanol, followed by vaporization of ethanol. Filter paper was inserted into a slit on foamed polystyrene (see Extended Data Fig. 1a). A larva was put on the edge of the slit and videotaped. All larvae palpated on filter paper (Extended Data Fig. 1b). The fraction of larvae showing test or persistent biting was investigated for groups of ten individuals. Statistical analysis was performed using Kruskal-Wallis test followed by Dunn’s test (Prism ver.5, Graph-Pad, La Jolla, USA).

### Agar-based food intake assay

To determine whether phytochemicals induce persistent biting, we used agar-based food intake assay^16^ with some modification. Agar-based diets basically contain 9% cellulose and 1% agarose in Elix water (Merck Millipore, Tokyo, Japan). Stimulants except for β-sitosterol and cellulose were mixed in advance and added to an agarose solution boiled using the microwave for dissolution. This mixture solution was vortexed well and poured into 90 mm petri-dish (Kanto chemical Co., Inc., Tokyo, Japan). After agar-based diet solidified, β-sitosterol dissolved in EtOH was applied to the diet followed by EtOH evaporation. For preparation of agar-based diets containing mulberry leaf extract, mulberry leaves were cut into small pieces and dried in a drying machine. Dried leaves were crushed into powder. The composition of the mulberry leaf extract diet was 75% mulberry leaf powder (w/v), 1% agarose (w/v), and 24% Elix water (v/v) based on ratio of dry and wet weight of intact mulberry leaves (data not shown). Three to five larvae were put on each petri-dish and larval weights before and after 3h feeding were measured. Increases of larval weight were regarded as amounts of food intake because no feces were observed after 3h feeding at 25°C. We adopted increases of body weight (mg) as the index of food intake because no significant correlation between amount of food intake and the original body weight was observed (r = 0.11, P > 0.5) by Pearson’s Correlation Analysis (Extended Data Fig. 3a). Statistical analysis was performed using Kruskal-Wallis test followed by Dunn’s test or two-way ANOVA followed by Turkey-Kramer test (Prism ver.5, Graph-Pad, La Jolla, USA).

### Tip recording

Tip recordings from all sensilla of MP and lateral and medial sensillum of MG were conducted to determine whether neurons respond to CGA, ISQ, Q3R and βsito. All eight sensilla in MP (Extended Data Fig. 2a) were stimulated together. LS and MS in MG have a *myo*-inositol neuron and a deterrent neuron that detects nicotine, respectively^13,14^, thereby they were used as positive control for stimulants of LS and MS (Extended Data Fig. 2b, 2c). Fifth instar larvae starved until use were used for recording. CGA, ISQ, and Q3R were dissolved and serially-diluted in water whereas βsito in ethanol. In dilution, solutions were stirred well for more than 30 min. We used 5 mM NaCl as an electrolytic solution. Methyl-β-cyclodextrin (βCD) was used as a complexing agent as described in Brown et al.^20^ with some modifications. A solution of 2 mg/mL βsito in ethanol was added into 2.5% βCD in 5% NaCl solution so that the final concentration of βsito was 50 μM. The solution was put on a heating block at 80°C until being clear. Tip recording methods were conducted based on Sasaki et al.^16^ Stimulants were filled up in glass microelectrodes and a silver wire inside of the electrode was connected to a TastePROBE amplifier (Syntech, Kirchzarten, Germany). The sensilla of MP and the sensilla of MG were capped with the recording electrode using a micromanipulator to stimulate the chemosensory neurons and to record the response simultaneously. As a reference electrode, a fine silver wire insulated to its tip was inserted into the base of the maxilla. Electrical signals were recorded on a computer via a PowerLab 4/25 and analyzed using CHART 5 (ADInstrument, Australia).

### Statistics

All statistical analyses were performed using Prism ver.8 (Graph-Pad, La Jolla, USA). We used parametric tests in Figs. 2a-d and 3b, c and Extended Data Fig. 1c, and non-parametric tests in Fig. 4a-c and Extended Data Fig. 3c, d. See Supplemental information for more details (*e.g.* exact sample sizes and P value).

## Data Availability

All datasets generated are available from the corresponding authors upon reasonable request.

