## Supplementary figures for "The two-step chemosensory system underlying the oligophagy of silkworm larvae"

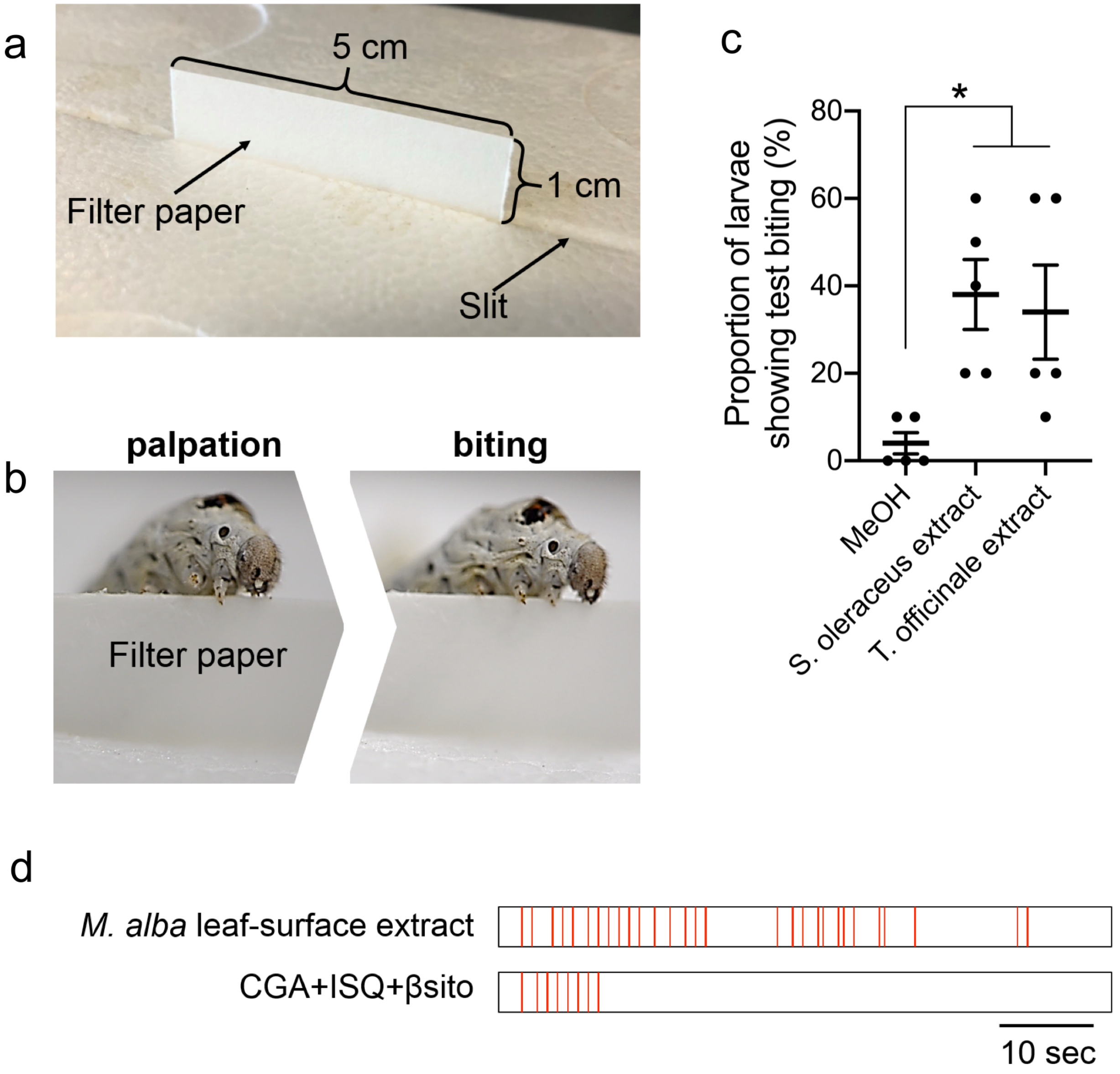


**Extended Data Figure 1 | Biting assay using filter paper. a,** Biting assay using filter paper. Filter paper was inserted into a slit on foamed polystyrene. **b,** All larvae palpated at the edge of the filter paper irrespective of treatment (left) and larvae showed test biting of treated filter paper. **c,** Proportion of larvae (n = 10) showing test biting of filter paper treated with leaf-surface extracts of two edible leaves of *S. oleraceus* and *T. officinale* over 1 minute. Experiments were repeated as independent biological replicates (n = 3–5). Data are means ± SE. Statistical analysis was performed using one-way analysis of variance (ANOVA) followed by Tukey *post hoc* test. An asterisk indicates a significant difference (**P* < 0.05). **d,** Representative raster plots of the timing and duration of biting behavior by larvae using filter paper treated with *M. alba* leaf-surface extract or a mixture of CGA, ISQ, and βsito.

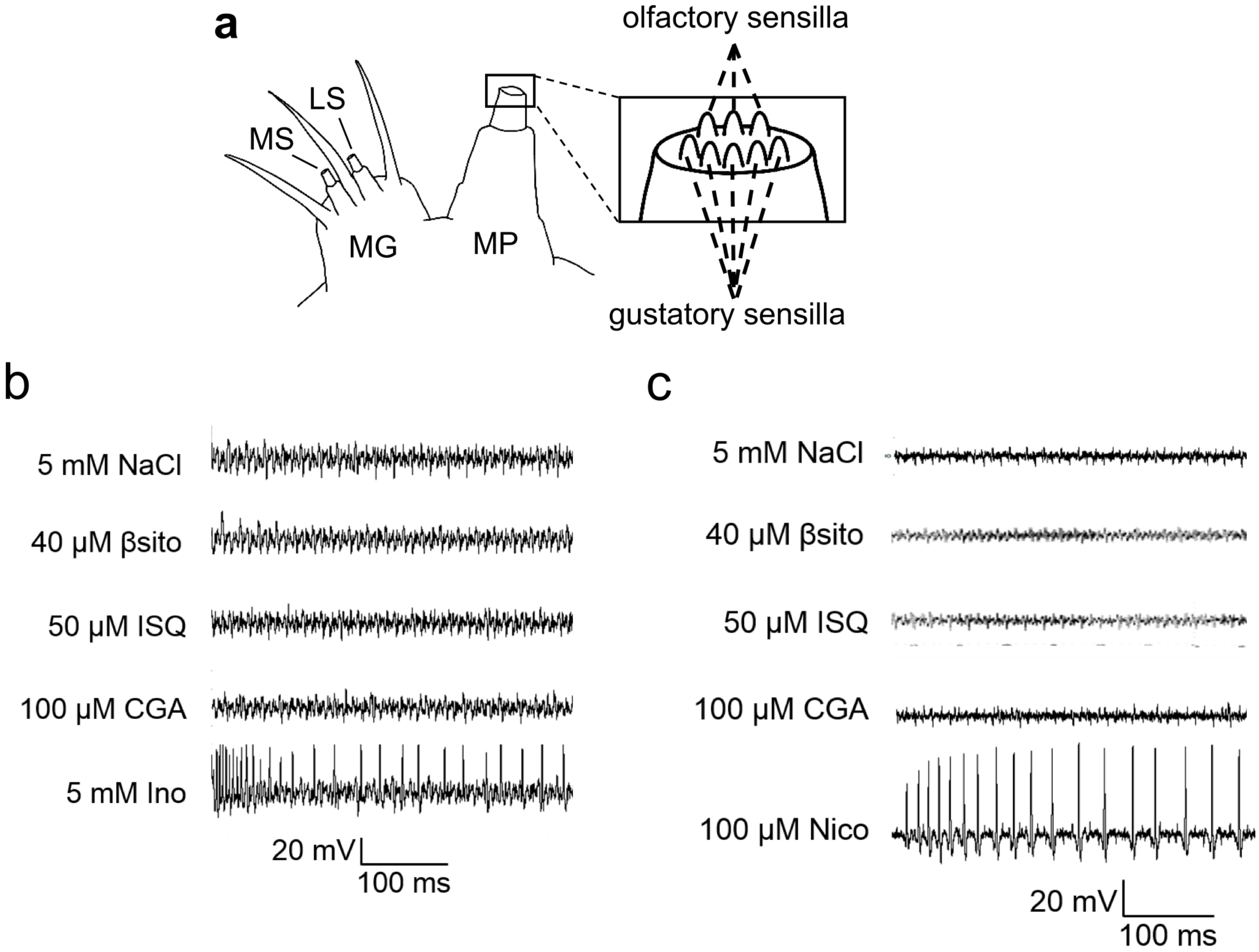


**Extended Data Figure 2 | Tip recording of sensilla in the MG. a,** Schematic of sensilla in the MP, which has eight sensilla (five putative gustatory and three olfactory sensilla). **b, c,** Typical electrophysiological recordings from LS **(b)** and MS **(c)** sensilla in the MG in response to chlorogenic acid (CGA), isoquercitrin (ISQ), quercetin-3-*O*-rhamnoside (Q3R), and β-sitosterol (βsito). *myo*-inositol (Ino) and nicotine (Nico) were used as positive controls for LS and MS sensilla, respectively.

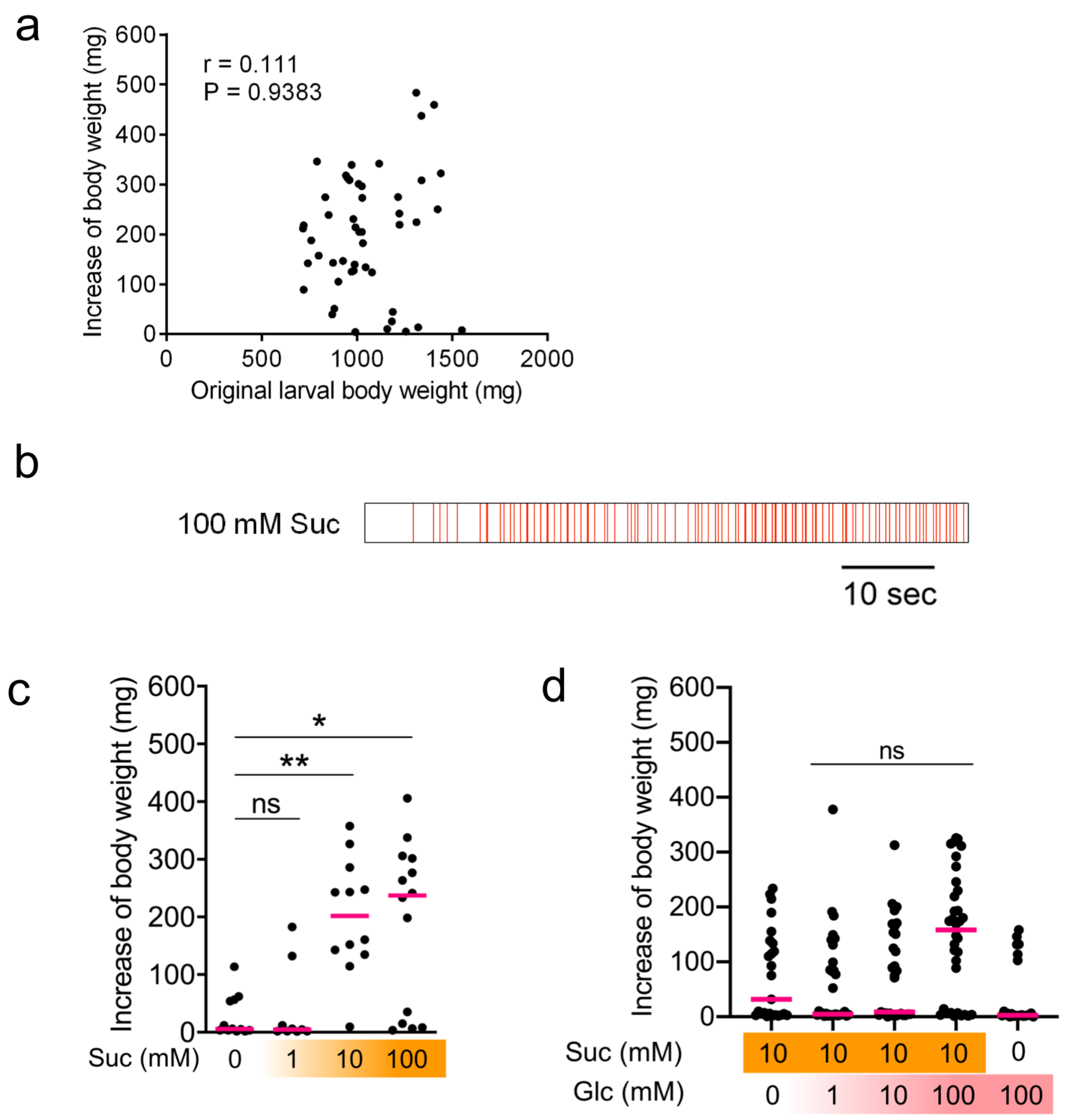


**Extended Data Figure 3 | Supplemental agar-based food-intake assay. a,** Correlation between the original larval body weight and the increase in body weight after feeding a 100 mM sucrose-containing agar-based diet for 3 hours. Correlation coefficient (r) by Pearson correlation analysis (n = 51). **b,** Representative raster plot of the timing and duration of biting when feeding agar containing 100 mM sucrose. **c,** Sucrose-dependent increase in larval weight in MP-ablated larvae after 3 hours. Magenta bars denote median. Statistical analysis was performed using Kruskal–Wallis test followed by Dunn test. “ns” indicates no significant difference; an asterisk indicates a significant difference (*$P$ < 0.05; **$P$ < 0.01).

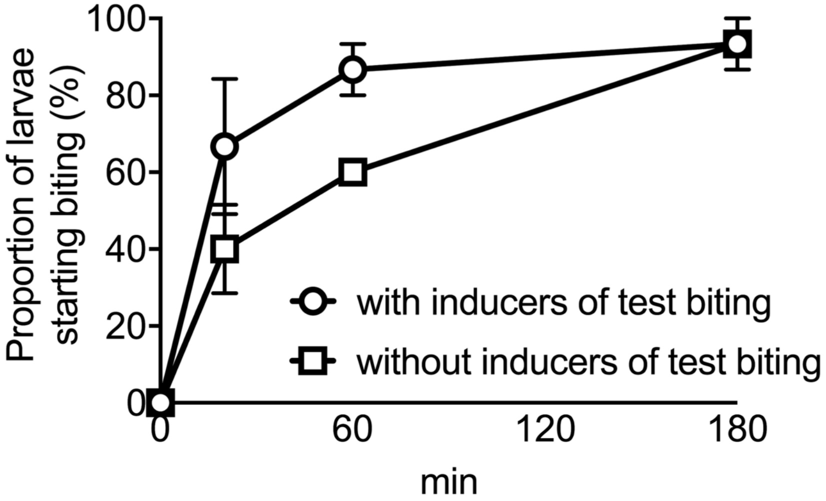


**Extended Data Figure 4 | Effect of a mixture of inducers of test biting (CGA, ISQ, and βsito) on increase in proportion of feeding larvae.** Sugars (10 mM sucrose, 5 mM *myo*-inostiol, and 5 mM glucose) were added to the basic agar food (9% cellulose and 1% agar). The following inducers of test biting were added: 100 μM CGA, 1 μM ISQ, and 3 μg/cm$^2$ βsito. Data are from biological triplicate experiments (n = 5); error bars indicate SE.
